# Functional composition has stronger impact than species richness on carbon gain and allocation in experimental grasslands

**DOI:** 10.1101/418301

**Authors:** Christiane Roscher, Stefan Karlowsky, Alexandru Milcu, Arthur Gessler, Dörte Bachmann, Annette Jesch, Markus Lange, Perla Mellado-Vázquez, Tanja Strecker, Damien Landais, Olivier Ravel, Nina Buchmann, Jacques Roy, Gerd Gleixner

**Author notes:** Corresponding author (GG).

## Abstract

Numerous experiments have shown positive diversity effects on plant productivity, but little is known about related processes of carbon gain and allocation. We investigated these processes in a controlled environment (Montpellier European Ecotron) applying a continuous ^13^CO_2_ label for three weeks to 12 soil-vegetation monoliths originating from a grassland biodiversity experiment (Jena Experiment) and representing two diversity levels (4 and 16 sown species). Plant species richness did not affect community- and species-level ^13^C abundances neither in total biomass nor in non-structural carbohydrates (NSC). Community-level ^13^C excess tended to be higher in the 16-species than in the 4-species mixtures. Community-level ^13^C excess was positively related to canopy leaf nitrogen (N), i.e. leaf N per unit soil surface. At the species level shoot ^13^C abundances varied among plant functional groups and were larger in legumes and tall herbs than in grasses and small herbs and correlated positively with traits as leaf N concentrations, stomatal conductance and shoot height. The ^13^C abundances in NSC were larger in transport sugars (sucrose, raffinose-family oligosaccharides) than in free glucose, fructose and compounds of the storage pool (starch) suggesting that newly assimilated carbon is to a small portion allocated to storage. Our results emphasize that the functional composition of communities is key in explaining carbon assimilation in grasslands.

## Introduction

Over the last years, numerous studies have accumulated evidence that loss of plant species diversity impairs ecosystem functioning [1]. While there is a growing consensus about positive relationships between plant species richness and primary productivity above- and belowground (e.g. [2,3]), the underlying physiological mechanisms are not well understood. Moreover, it has also been verified that the effects of increasing plant diversity on the biomass production of individual species are highly variable and range from positive, over neutral to negative effects (e.g. [4,5]). Photosynthetic carbon assimilation as well as the allocation and partitioning of carbon within the plant (as determined by source-sink relationships) are intimately linked to plant growth and productivity. Therefore information on how plant diversity affects these processes could contribute to a mechanistic understanding of diversity-productivity relationships and the varying responses of individual species to plant diversity.

Light and nitrogen availability are the primary limiting resources constraining carbon gain in temperate grasslands under humid climate. Photosynthetic enzymes contain a major portion of nitrogen (N) in leaves and leaves with higher N contents have in general higher rates of photosynthesis [6]. Canopy photosynthesis depends on the distribution of N among the leaves as well as on the total amount of leaf area (i.e. LAI; leaf area index) of the stand [7]. Biodiversity experiments in temperate grasslands have shown that canopy height, density and LAI increase with species richness [8,9]. Grassland vegetation usually consists of a mixture of species of various heights. Incoming photosynthetic active radiation decreases with increasing depth in the canopy [10]. The vertical light gradient goes along with changes in leaf photosynthetic capacity, which decreases with lower light availability in the lower canopy. Leaves in different positions of the canopy have different physiological potentials, which are finely tuned to the prevalent light conditions [11]. This acclimation in photosynthetic capacity involves variation in a range of leaf traits between the bottom and the top of the canopy, such as leaf N concentrations, specific leaf area or stomatal conductance [12-14]. In addition, the structural changes of vegetation entail variation in other microclimate factors such as temperature, air humidity and intra-canopy CO_2_ concentrations [15,16], which also sensitively interact with photosynthetic gas exchange.

In leaves, synthesis of carbohydrates (sucrose, starch) and N assimilation (amino acid synthesis) are the major sinks for products of photosynthesis. During the day, new assimilates are partitioned between sucrose for immediate demands of the plants on the one hand, and starch for growth-related demands during the night on the other [17]. The export of carbohydrates (mainly as sucrose) from photosynthetically active “source” tissue such as mature leaves provides the base for growth and maintenance of non-photosynthetic “sink” tissues (roots, plant organs related to reproduction) or those with higher demands of carbon supply (growing tissues). Carbon supply to sinks goes either as direct transfer of products of the current photosynthesis or diurnal or longer-term reserve pools are reallocated to the sink. Reserve pools are important for carbon supply in the dark of diurnal cycles, but they can also support sinks during periods with active photosynthesis [18]. There are only few studies that assessed the partitioning of ^13^C label originating from ^13^CO_2_ between different non-structural carbohydrates (NSC) (e.g. [19,20]). When sink activity is not reduced by drought or other stressors, a relative larger amount of new assimilates seems to be allocated to sugars compared to storage compounds [21]. So far, it is not known if differences in plant diversity impact the allocation of carbon into different plant compartments (root, shoots) or into carbohydrates with different functions (such as transport or storage).

In mixed plant communities, the degree to which species with different combinations of physiological and morphological traits are able to acquire essential resources and use these resources for growth determines community-level carbon gain [22]. Community-level metrics derived from functional traits are increasingly adopted as predictors for ecosystem functioning. Community-weighted trait means (CWM; [23]) and functional trait diversity based on the Rao coefficient (FD; [24]) describe two complementary aspects of the relationship between the functional composition and ecosystem functioning [25]. CWMs conceptually follow the assumption that ecosystem processes are mainly determined by trait values of the dominant species. FD quantifies the dispersion of trait values in a community and has been linked to niche complementarity assuming that a greater variety of trait values reduces niche overlap among species [26], but to date trait-based metrics were mostly related to community biomass as a measure of plant productivity in biodiversity experiments (but see [27]).

Photosynthate labelling methods using isotopically enriched ^13^CO_2_ may help to estimate the amount of carbon dioxide taken up for assimilation and the allocation and distribution of carbon in the plant (e.g. [28,29]). In particular, steady-state labelling (i.e. continuous ^13^C labelling of all assimilated CO_2_ at constant enrichment level for a longer duration), which even allows slow turn-over pools to reach steady state, might be useful for estimating carbon gain of all species in a mixed community. However, continuous labelling of mixed plant communities in the field is challenging and has never been applied in a biodiversity experiment. Here, we capitalized on transferring twelve large soil-vegetation monoliths originating from two levels of sown diversity (4 and 16 species) from a long-term biodiversity experiment (Jena Experiment [30]) into an advanced controlled environment facility (Montpellier European Ecotron [27]). The monoliths were treated with a continuous ^13^CO_2_ label over a period of three weeks followed by immediate harvest of the plants to trace the fate of the labelled C in bulk shoot and root biomass as well as in NSC, which are known to have different functions (e.g. transport and storage) in the plant’s carbon metabolism.

In a previous study we have shown that gross and net ecosystem C uptake rates were higher at increased species richness and that higher leaf area index (LAI) and diversity in leaf nitrogen concentrations were most important in explaining positive diversity effects [27]. In this study, however, it was not possible to directly compare the diversity effects at community level versus species-level responses or the allocation of carbon into different plant compartments and into carbohydrates with different functions. Therefore, the specific goals of the ^13^C labelling study were firstly to test how plant diversity affects carbon gain at community and species level, as well as the allocation of carbon from shoots to the roots and into different NSC. The second key aspect of our study was to evaluate which plant traits related to leaf gas exchange, photosynthetic capacity and plant positioning in the canopy and which metrics describing their distribution in the plant communities may explain differences in species- and community-level carbon gain. We tested the following hypotheses: (1) The amount of assimilated ^13^C (i.e. ^13^C excess) increases with species richness of the communities due to greater community biomass and LAI. (2) The ^13^C abundances in shoots are not dependent on species richness of the communities but vary with leaf nitrogen concentrations of the involved species. (3) The ^13^C abundances in different NSC are higher in carbohydrates related to transport than in those related to storage irrespective of community species richness but depend on species traits related to photosynthesis and positioning in the canopy.

## Material and methods

### Plant communities originating from the Jena Experiment

Plant communities originated from the Jena Experiment, a long-term grassland biodiversity experiment located in the floodplain of the river Saale (Jena, Germany, 50°55’N, 11°35’E,130 m a.s.l.; [30]). The experimental area was an agricultural field before different combinations from a pool of 60 species typical for Central European mesophilic grasslands (*Arrhenatherion* community [31]) were sown in spring 2002. Briefly, the Jena Experiment consists of 82 large plots (20 × 20 m size) representing all possible combinations of six species-richness levels (1, 2, 4, 8, 16, and 60) crossed with four levels of functional group richness (1 to 4; legumes, grasses, small herbs, tall herbs). Plots are arranged in four blocks to account for a gradient in soil texture from sandy loam to silty clay with increasing distance to the riverside. Plots are mown twice per year (early June, September) and mown biomass is removed to mimic the typical management of hay meadows in the region. Plots do not receive any fertilization. All plots are regularly weeded (at the beginning of the growing season and after first mowing) to remove unwanted species.

Twelve plots representing two sown species-richness levels (4 and 16 species) were chosen according to the following criteria: (1) mixtures contained legumes, grasses and herbs, (2) realized species numbers were close to sown species richness, and (3) plots were equally distributed among the blocks of the Jena Experiment. The selected plots met the above-mentioned criteria with the exception of one 4-species mixture, which did not contain grass species (see S1 Table for species compositions).

In December 2011, soil monoliths (2 m depth, 1.6 m diameter) selected to be representative for the plots of origin (as percentage vegetation cover) were excavated using steel lysimeters (UMS GMBH, Munich, Germany) (see also [32]). After extraction lysimeters were buried to soil surface level at the Jena Experiment field site being exposed to the same environmental conditions as the plots.

### Experimental set-up in the CNRS Ecotron facility

The lysimeters were transported to the CNRS Ecotron facility in Montpellier (France) at the end of March 2012. There, the monoliths were randomly allocated to the 12 experimental units of the macrocosm platform [27]. The belowground compartment was maintained in the lysimeter. The aboveground compartment was installed in controlled environment units (30 m^3^ domes) exposed to natural light conditions and UV radiation retained by the cover with 250 μm thick Teflon-FEP film (DuPont, USA). The main abiotic characteristics of the ecosystem atmospheric compartment (air temperature, humidity, CO_2_ concentration) were controlled in each unit. The monoliths were kept under controlled conditions for four months, spanning the phase of high vegetative growth (end of March to end of July) before the experiment was finished with a destructive harvest. The imposed climate regime intended to simulate the average climatic conditions at the Jena Experiment field site since the start of the biodiversity experiment in 2002. As the temperature and precipitation regime in spring/summer 2007 were very close to average conditions, the 10 min-interval data from the Jena Experiment weather station containing the daily profiles of air temperature and humidity were imposed as climatic set points in the Ecotron (for details see [27]). To balance out differences in the incoming radiation between Jena and Montpellier (37% lower in Jena during the period from April to July), a black shading mesh was installed on the inside of each dome, which reduced the incoming radiation by 44%.

The monoliths were weeded twice during the experiment period (April, May) to maintain the target species composition. Shoot biomass was mown at the end of April to mimic the mowing management of the Jena Experiment.

### ^13^CO_2_ labelling of the experimental units

A continuous atmospheric ^13^CO_2_ labelling was applied during the day in all units for a period of three weeks (4-24 July 2012) using an automated system. A cylinder of compressed 99% ^13^CO_2_ (Eurisotop, France) was connected to a manometer and a high accuracy mass flow controller (F200CV, Bronkhorst, Netherlands). The ^13^CO_2_ was injected in each unit as 30 s pulses every 6 min. During a pulse, the flow rate of ^13^CO_2_ was regulated at 4.8 ml min^-1^ and injected along with roughly 0.5 l min^-1^ of CO_2_-free air, using a Valco valve (EUTA-SD16MWE dead-end path, VICI, USA). The labelling system was operated and data were logged using a PXI Chassis (National Instrument, USA), a modbus with a RS232 communication module and LabVIEW programming (National instrument, USA). The δ^13^C of CO_2_ at the outlet of each unit was monitored on-line every two hours using a Picarro G2101-i Isotope Analyzer (Picarro, USA), and an in-house automatic manifold. The obtained δ^13^C-CO_2_ value was +21.21‰ (mean across 12 experimental units, SD = 2.23‰).

### Sampling of shoot and root biomass

Aboveground biomass was sampled by cutting the plant material 3 cm above ground-level in two randomly positioned rectangles (each 50 × 20 cm) in late April (30 April – 1 May 2012) before labelling in parallel to mowing of the experimental units (see above). Each sample was sorted by target species (= sown into a particular community), weeds and detached dead material. Because not all target species growing in an experimental unit were present in the biomass samples, additional plant material for stable isotope analyses of missing species was sampled by harvesting three to five shoots if possible from different individuals in the remaining area. All samples were dried at 65°C for 48h and weighed.

Aboveground plant material of each present species per experimental unit was sampled again at the end of the ^13^CO_2_ labelling period (20 July 2012) for stable isotope analyses. Three to five shoots randomly chosen on half of the unit, which was available for sampling, were cut near ground-level and dried at 65°C for 48h. Shoot biomass production was determined from a final destructive harvest of the Ecotron experiment by clipping the vegetation at ground-level on an area of 90 × 55 cm in each unit (23-24 July 2012). Samples were sorted by species, and a subsample per species and experimental unit was separated into compartments (leaf, stems, reproductive organs). All samples were dried at 65°C for 48h and weighed (see S1 Table for species present in biomass samples of each experimental unit).

Root biomass was sampled by collecting three cores (inner Ø 3.5 cm) down to a depth of 60 cm in each experimental unit (17 July 2012). Cores were divided in depth increments (0-5, 5-10, 10-20, 20-30 and 40-60 cm) and sections of the same depth were pooled per experimental unit. Samples were washed with tap water over a sieve of 200 μm mesh size to separate roots from soil. Root biomass was dried at 65°C for 48 hours before weighing and calculating root biomass per soil volume (mg_root_ cm^-3^_soil_).

### δ^13^C analysis and isotope calculations

Shoot samples per species and experimental unit from both harvests and root samples per depth increment were analysed with an isotope ratio mass-spectrometer (Delta C prototype Finnigan MAT, Bremen, Germany). Our calculation to determine ^13^C enrichment and ^13^C excess after the labelling period (July) in relation to the background values before the labelling were based on values of δ^13^C, which express the ratios of ^13^C/^12^C in the sample (R_sample_) over the ^13^C/^12^C ratio of a standard R_standard_ (V-PDB for carbon = 0.0111802) in ‰,

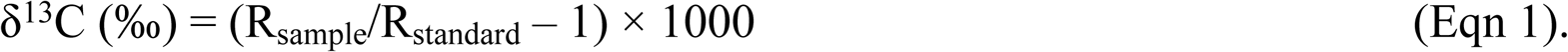

Background values for shoot-level data were taken from species-specific shoot δ^13^C measured for each unit from the harvest in April before the labelling. Because background values for roots were not available, the proportion of each species in the aboveground biomass of the final harvest in July and the respective shoot δ^13^C background values were used to derive biomass-weighted δ^13^C values for each unit. To account for the modification of the isotopic signature of different plant organs through post-photosynthetic fractionation, which has been reported to result in a general ^13^C enrichment of roots compared to shoots (∼0.96 ‰ according to meta-analysis across 88 studies; [33]), we assumed an offset of 1 ‰ to the biomass-weighted community-level δ^13^C values to obtain background values for the roots.

The δ^13^C values (‰) of background samples and labelled samples were back-converted into absolute isotope ratios according to Eqn 1. Then, R_sample_ was related to fractional abundance (A)

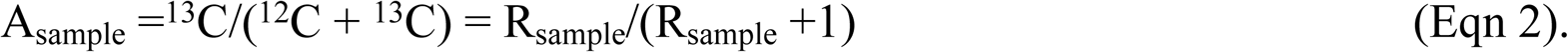

The differences between the fractional abundances of labelled and background samples served to assess ^13^C atom% excess.

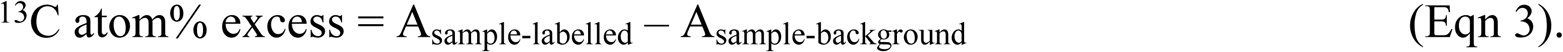

The ^13^C abundance (µg g_dw_ ^-1^) was obtained as

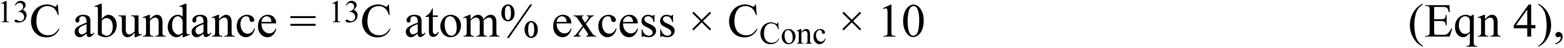

where C_conc_ are tissue carbon concentrations (mg C g_dw_^-1^). Species-level (shoot biomass) and depth-level (root biomass) were weighted by biomass of the species or depth level, respectively, to get shoot and root community-level ^13^C abundances.

The fractional abundances of the labelled samples and carbon content (= C_sample_; mg m^-2^) derived from C_conc_ (mg C g_dw_ ^-1^) and biomass (g_dw_ m^-2^) were used to calculate ^13^C excess (mg m^-2^) as

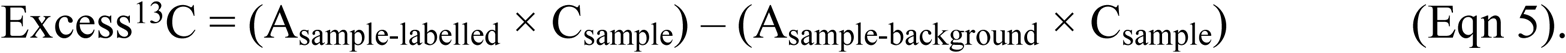

Excess^13^C per species (shoot biomass) and depth layer were summed to derive community-level excess ^13^C.

### Analysis of non-structural carbohydrate concentrations and δ^13^C

Three additional soil cores (inner Ø = 4.8 cm) were collected using a split-tube corer (Eijkelkamp Agrisearch Equipment, Giesbeek, The Netherlands) from each experimental unit (16-18 July 2012; 10 a.m.-2 p.m.) for analyses of NSC. Shoot material was cut-off from each core. Afterwards, soil cores were pooled per experimental unit. Fresh soil samples were immediately sieved (2 mm mesh size) and all roots were carefully collected. Shoot and root material from soil cores from each experimental unit were placed in separate bags, shock-frozen in liquid nitrogen and subsequently freeze-dried (for details see [34]).

Samples of 3 to 12 fully-expanded leaves (dependent on leaf size) were collected in each experimental unit at midday (1-2 p.m.) shortly before the end of the ^13^CO_2_ labelling period (on 20 July 2012). In total 34 samples were taken from species, which were either available at both low and high diversity (samples of 6 species) or being well established representatives of each functional group (11 samples in 4-species mixtures, 23 samples in 16-species mixtures; see S1 Table for species identities). Samples were shock-frozen in liquid nitrogen and stored at –80°C in a freezer until vacuum freeze-drying.

A detailed description of the extraction and measurement methods for non-structural carbohydrates (NSC) is provided in the S1 Methods. Briefly, total water soluble sugars were extracted from freeze-dried and finely milled plant material using a hot water extraction method adapted from [35]. After removing charged compounds, the neutral fraction was analysed by high-performance liquid chromatography-isotope ratio mass spectrometry (HPLC-IRMS, ThermoFinnigan LC-IsoLink System and Deltaplus XP IRMS, Thermo Electron, Bremen, Germany) using a Nucleogel Sugar 810 Ca^2+^column (Macherey & Nagel, Düren, Germany). This method provided a good separation of sucrose, glucose and fructose. Peaks of oligosaccharides, with retention times corresponding to the external standards raffinose, stachyose and verbascose, overlapped and were integrated as one single peak. These sugars represent the same functional type (symplastic phloem transport and short-term storage) and were described as raffinose-family oligosaccharides (RFOs) [36,37]. However, it was not possible to distinguish RFOs from isomeric oligosaccharides, such as fructans, with the applied method. Mass spectrometry analyses and acidic hydrolysis of sugar extracts indicated the presence of both, RFOs and fructans.

Similar to soluble sugars, starch was extracted from freeze-dried and finely milled plant material using an enzymatic method adapted from [38,39]. First, soluble sugars were removed by incubating the samples with a methanol:chloroform:water mixture, then starch was hydrolysed with heat stable α-amylase. After removing the enzyme, starch hydrolysates were analysed on HPLC-IRMS (see above) without prior separation on a column.

### Vegetation-related predictor variables

Canopy leaf nitrogen (g N_Leaf_ m_Soil surface_^-2^) as a proxy for photosynthetic assimilation capacity was estimated as the sum of species-specific values for leaf area (m^-2^) × specific leaf mass (g_Leaf_ m_Leaf_ ^-2^) × leaf N concentration (g N g_Leaf_ ^-1^), where total leaf area per m^2^ was obtained from specific leaf area and leaf mass derived from separating subsamples per species into plant compartments during the final harvest (as described above).

Alternatively, aboveground plant traits expected to be relevant for carbon uptake and metabolism were measured for all available plant species in each experimental unit: specific leaf area (SLA), leaf dry matter content (LDMC), stomatal conductance (g_s_), leaf greenness (LeafG), leaf N concentrations (N_Leaf_), and shoot height (H_Shoot_). See S2 Methods for a detailed description of trait measurements. Trait values and species biomass proportions were used to derive community-weighted trait means [23] according to the equation

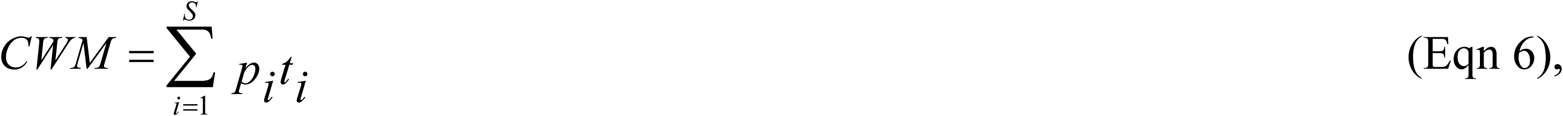

and trait diversity (FD) based on Rao’s quadratic entropy [24]

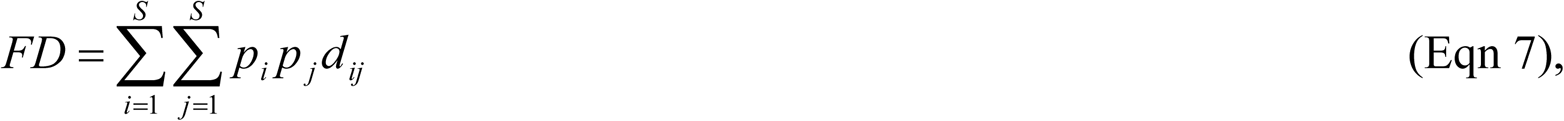

where *S* is the number of species in the community, *t*_*i*_ is the trait value of species *i, d*_*ij*_ is the trait distance between species *i* and *j*, and *p*_*i*_ and *p*_*j*_ are the biomass proportions of species *i* and *j* in the respective mixture. Calculations of trait-based indices were conducted using the R library *FD* [40].

### Statistical analyses

Data analysis was performed with the statistical software R3.0.2 [41]. Species richness effects on community-level ^13^C abundance and ^13^C excess in shoot and root biomass were tested with analysis of variance (ANOVA). To identify the most important predictors for each response variable, simple linear regressions were fitted for each potential predictor (i.e. canopy leaf N, as well as CWM and FD of single traits (N_Leaf_, LeafG, SLA, LDMC, g_s_, H_Shoot_). Other data were analysed with linear mixed-effects models using the *lmer* function implemented in the *lme4* library [42]. For community-level ^13^C abundance in bulk shoot and root NSC, analysis started from a constant null model with mixture (= experimental unit) as random effect. Species richness (SR; 4 species vs. 16 species) and terms for the identity of non-structural carbohydrates (NSC-ID; five factor levels: glucose, fructose, sucrose, RFO, starch) and plant compartment (two factor levels: shoot, root) as well as the two- and three-way interactions between these terms were sequentially added. For species-level data, mixture (= experimental unit) and plant species identity (Species-ID) were fitted as independent random effects to account for multiple measurements in the same mixture and the same species in different mixtures. In the model for species-level ^13^C abundance in shoot biomass, species richness and functional group identity (FG-ID; four factor levels; legumes, small herb, tall herbs, grasses) and the interaction between both terms were fitted as fixed effects. In analyses of leaf carbohydrates additional terms for the identity of the carbohydrates and the respective two- and three-way interactions with SR and FG-ID were entered. The maximum likelihood method (ML) was applied and likelihood ratio tests (χ^2^ ratio) were used to compare succeeding models and test for a significant model improvement by the added fixed effects. To identify differences between particular functional groups or carbohydrates, Tukey’s test was applied using the *glht* function in the R library *multcomp* [43] in models fitted with restricted maximum likelihood (REML).

Afterwards, in species-level analyses traits considered to be relevant for carbon gain (N_leaf_, g_s_, SLA, LDMC, H_shoot_) were explored as predictor variables in mixed-effects models with the random structure as described above. First, alternative models were fitted with each predictor variable separately as fixed effect. Models were compared using Akaike’s information criterion (AIC), which measures the lack of the fit with respect to the complexity of a model [44]. The model with the lowest AIC containing one of the predictor variables was chosen and extended fitting in alternative models one of the remaining predictor variables. Again, the model with the lowest AIC was chosen and likelihood ratio tests were applied to compare the best extended model with its predecessor until additional terms did not led to a significant model improvement and the model could be considered as minimal adequate.

## Results

### Community-level ^13^C abundance and ^13^C excess in shoot and root biomass

The ^13^C abundance in community shoot or root biomass did not differ between the 4-species and the 16-species mixtures (Figs 1a, b). Community-level ^13^C excess, i.e. the total amount of ^13^C in plant biomass, tended to be higher in the 16-species than in the 4-species mixtures, but differences between both diversity levels were not significant (F_1,10_ = 4.13, P = 0.067). Community-level shoot or root ^13^C excess also did not differ significantly between both diversity levels (Figs 1d, e). On average, shoot ^13^C excess was three times larger than root ^13^C excess (Fig 1f). This was mainly due to higher ^13^C abundances in shoots than in roots (on average 18 times greater; Fig 1c) since shoot biomass was considerably smaller than standing root biomass (shoot:root ratio 0.193 ± 0.057 SD; Figs 1g-i) and it was the entry point of the ^13^CO_2_ label.

**Fig 1.**
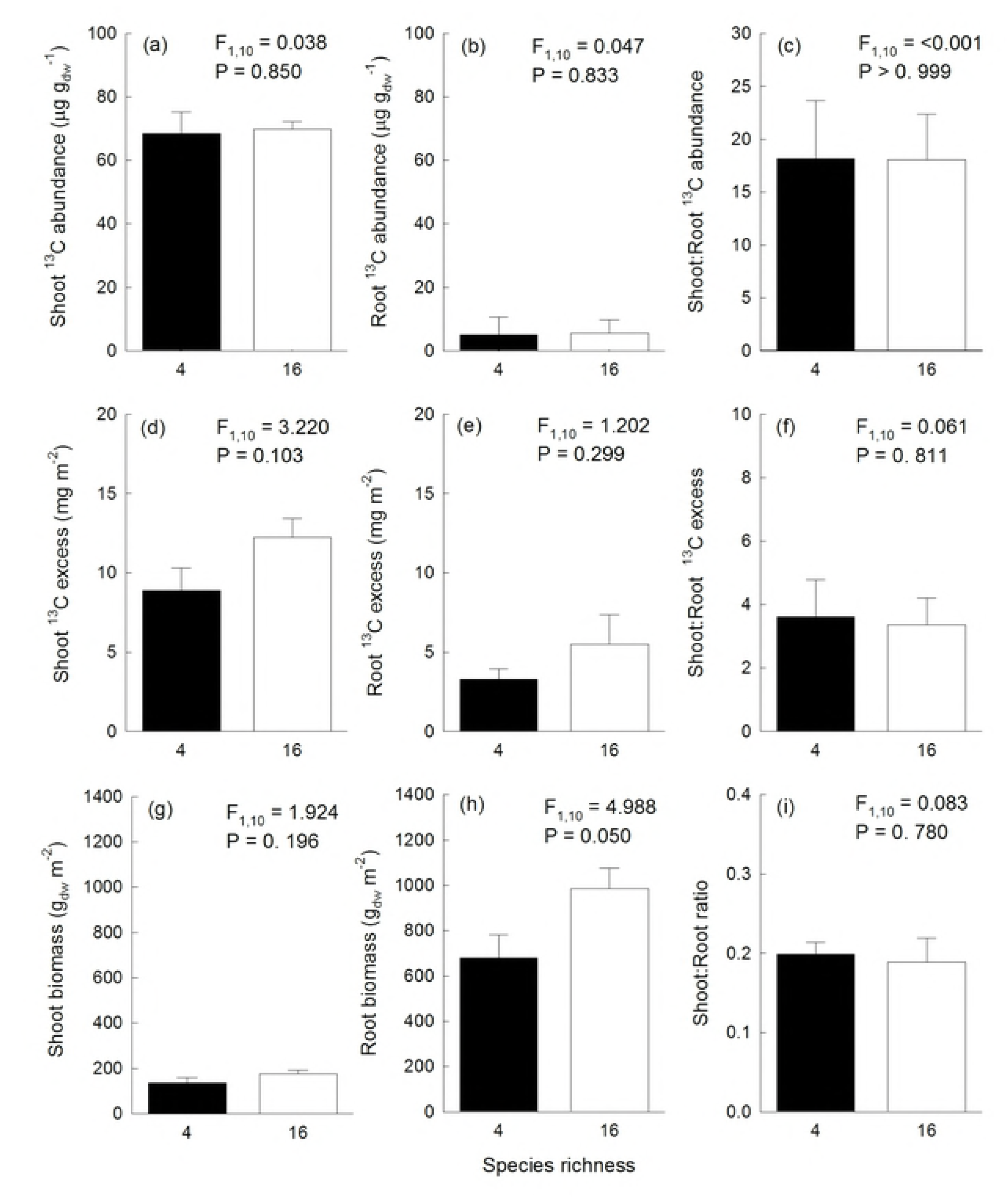
Community-level ^13^C abundance, ^13^C excess and biomass in relation to species richness. Relationships between species richness and community-level (a) ^13^C abundance in shoot biomass, (b) ^13^C abundance in root biomass, (c) shoot:root ratio of ^13^C abundances, (d) shoot ^13^C excess, (e) root ^13^C excess, (f) shoot:root ratio of ^13^C excess, (g) shoot biomass, (h) root biomass, and (i) biomass shoot:root ratio. F statistics and P values were obtained from ANOVA testing for differences between diversity-levels.

Community-level ^13^C abundance in shoot biomass was positively related to community-weighted means in leaf N (CWM-N_Leaf_; Fig 2a). Community-weighted means (CWM) or diversity (FD) in other studied traits did not show significant relationships with ^13^C abundance in shoots or community-level shoot ^13^C excess (S2 Table). However, community-level shoot ^13^C excess increased with increasing canopy leaf N, i.e. the amount of leaf N per unit soil surface (Fig 2b). Among the terms determining canopy leaf N, i.e. total leaf area, community-weighted means in SLA and N_Leaf_, the relative importance (expressed as the R^2^ contribution averaged over ordering among regressors) of total leaf area (lmg = 0.951) was greatest compared to minor relative importance of CWM-N_Leaf_ (lmg = 0.015) and CWM-SLA (lmg = 0.034).

**Fig 2.**
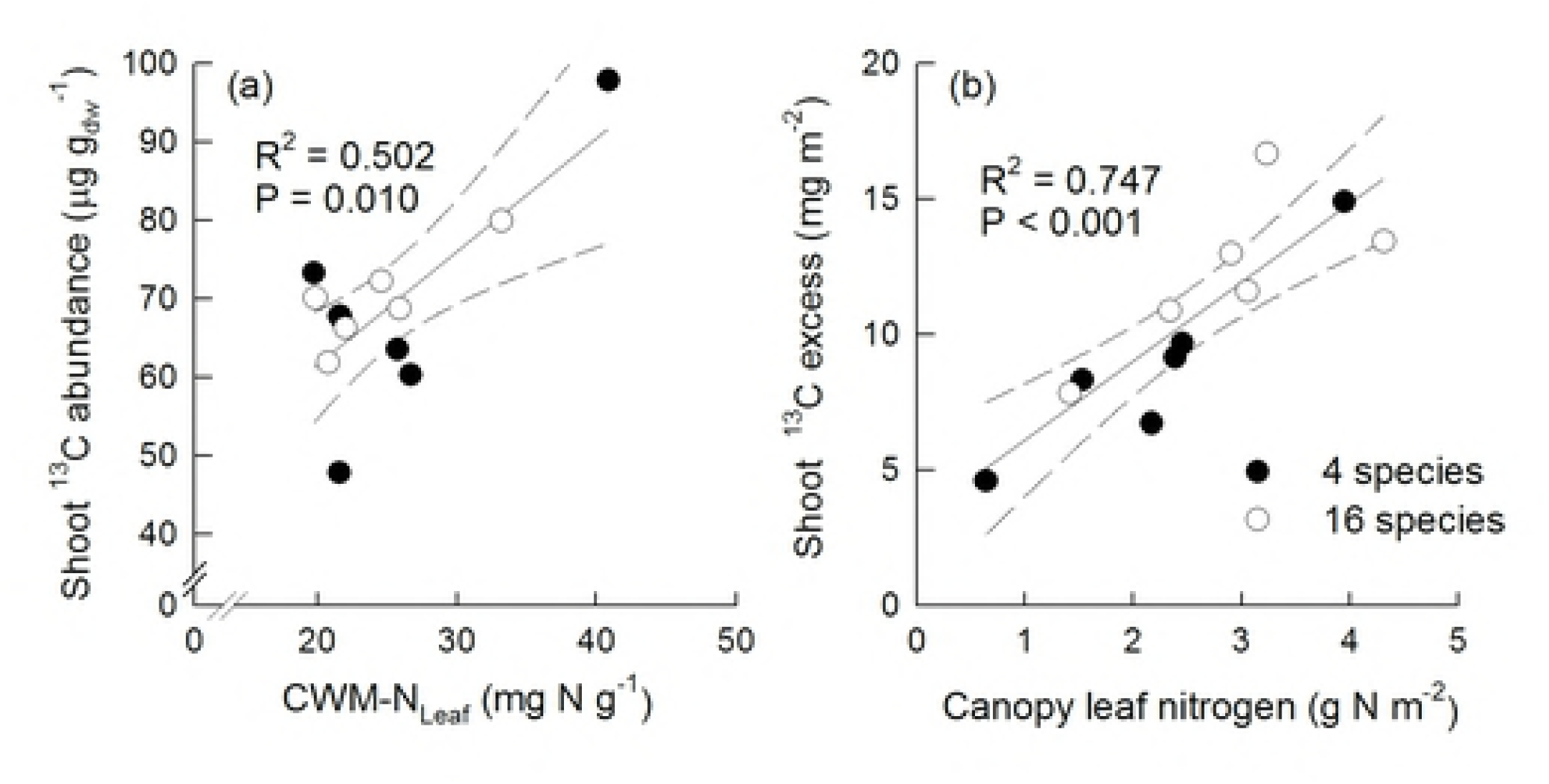
Relationships between community-level ^13^C abundance in shoot biomass and biomass-weighted means of leaf nitrogen concentrations (a), and relationships between community-level shoot ^13^C excess and canopy leaf nitrogen per unit soil surface (b). Closed circles display mixtures sown with 4 species, open circles display mixtures sown with 16 species.

### Species-level ^13^C abundances in shoot biomass

Species-level ^13^C abundances in shoot biomass were not different between mixtures with 4 and 16 sown species (χ^2^ = 1.600, P = 0.206). Species-level ^13^C abundances differed among functional groups (χ^2^ = 14.845, P = 0.002). Functional-group differences did not depend on diversity (non-significant interaction SR × FG-ID; χ^2^ = 1.089, P = 0.780). On average, legumes had higher ^13^C abundances per unit dry mass than species of the non-legumes functional groups, especially grasses and small herbs, while differences between tall herbs and legumes were statistically not significant (Fig 3). Species-level ^13^C abundances were positively related with N_Leaf_ (χ^2^ = 14.40, P < 0.001), g_s_ (χ^2^ = 5.75, P = 0.016), LeafG (χ^2^ = 5.48, P = 0.019) and H_Shoot_ (χ^2^ = 7.05, P = 0.008) (S1 Fig). The model best explaining variation in species-level ^13^C abundances included positive effects of N_Leaf_, g_s_ and H_Shoot_ (Table 1).

**Table 1.** Analysis of species-level ^13^C abundances in shoot biomass. Shown is the summary of a mixed-effects model testing for relationships between species-level ^13^C abundances in shoot biomass and plant traits.

| | $^{13}\text{C}$ abundance |
| --- | --- |
| AIC | 582.90 |
| **Fixed effects** |  |
| Intercept | -39.981 |
| $N_{\text{Leaf}}$ | 24.777 |
| $g_s$ | 10.985 |
| $H_{\text{shoot}}$ | 9.136 |
| **Random effects** |  |
| Unit | <0.001 |
| Species ID | 6.078 |
| Residuals | 14.772 |

**Fig 3.**
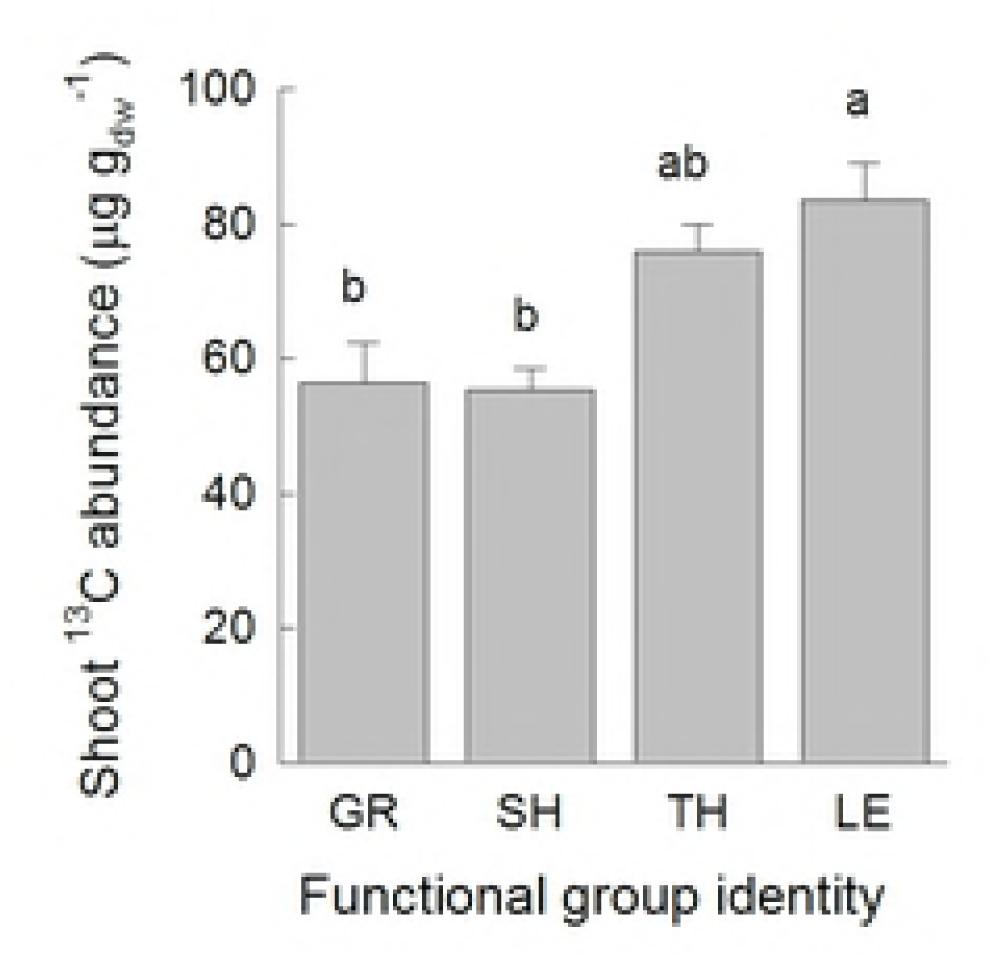
Species-level ^13^C abundances in shoot biomass. Shown are means (+1 SE) per functional group. Results of Tukey’s test (P < 0.05) applied to test for significant differences among functional groups are indicated with letters. Abbreviations are: GR = grasses, SH = small herbs, TH = tall herbs, LE = legumes.

The adequate set of predictor variables was determined by stepwise inclusion of predictor variables and model comparisons. Estimated coefficients are given for the resulting best model. Intercepts and slopes are shown for the fixed effects, and estimated standard deviations are given for the random effects. Abbreviations: g_s_ = stomatal conductance, H_Shoot_= shoot height, N_Leaf_ = leaf nitrogen concentration.

### ^13^C abundances in non-structural carbohydrates

The ^13^C abundance in NSC per unit dry mass in bulk shoot and root material did not differ between mixtures with 4 and 16 sown species (Table 2). The ^13^C abundance in NSC was higher in shoots than in roots for glucose, fructose, sucrose and starch (Fig 4a). Higher shoot-levels of ^13^C abundances in these NSC were attributable to both, greater ^13^C atom% excess and higher concentrations in shoots than in roots (Figs 4b, c). The ^13^C abundance in RFOs was not different between shoots and roots. Greater ^13^C atom% excess in RFOs in shoots was counterbalanced by higher concentrations of RFOs in roots. Overall, ^13^C abundances were larger in sucrose and RFOs than in glucose, fructose or starch (Fig 4a). The ^13^C atom% excess in roots and shoots was highest in sucrose and explained the ^13^C abundances in this metabolite. The ^13^C atom% excess in other NSC was lower, especially in the roots, and the lowest ^13^C atom% excess was found in starch stored in the roots. The high ^13^C abundance in RFOs was due to high concentrations especially in the roots.

**Table 2.** Analysis of community-level non-structural carbohydrate concentrations and δ^13^C. Shown is the summary of mixed-effects models testing for differences between plant species richness levels, non-structural carbohydrates (NSC-ID: glucose, fructose, sucrose, RFO, starch) and plant compartment (shoot, root) in ^13^C abundance, ^13^C atom% excess and concentrations.

| Source of variation | $^{13}\text{C}$ abundance | | $^{13}\text{C}$ atom% excess | | Concentration | |
| --- | --- | --- | --- | --- | --- | --- |
| | $\chi^2$ | P | $\chi^2$ | P | $\chi^2$ | P |
| Species richness (SR) | 0.86 | 0.354 | 1.24 | 0.266 | 0.07 | 0.797 |
| NSC-ID | 62.03 | <0.001 | 44.00 | <0.001 | 68.93 | <0.001 |
| SR x NSC-ID | 1.13 | 0.889 | 0.69 | 0.952 | 4.07 | 0.397 |
| Plant Compartment (Comp) | 51.98 | <0.001 | 100.68 | <0.001 | 13.80 | <0.001 |
| Comp x SR | 0.13 | 0.716 | 0.19 | 0.663 | 0.26 | 0.612 |
| Comp x NSC-ID | 14.65 | 0.005 | 34.18 | <0.001 | 34.60 | <0.001 |
| Comp x SR x NSC-ID | 0.11 | 0.998 | 0.56 | 0.968 | 4.13 | 0.389 |

Models were fitted by stepwise inclusion of fixed effects. Listed are the results of likelihood ratio tests ( $\chi^2$ ) that were applied to assess model improvement and the statistical significance of the fixed effects (P values).

**Fig 4.**
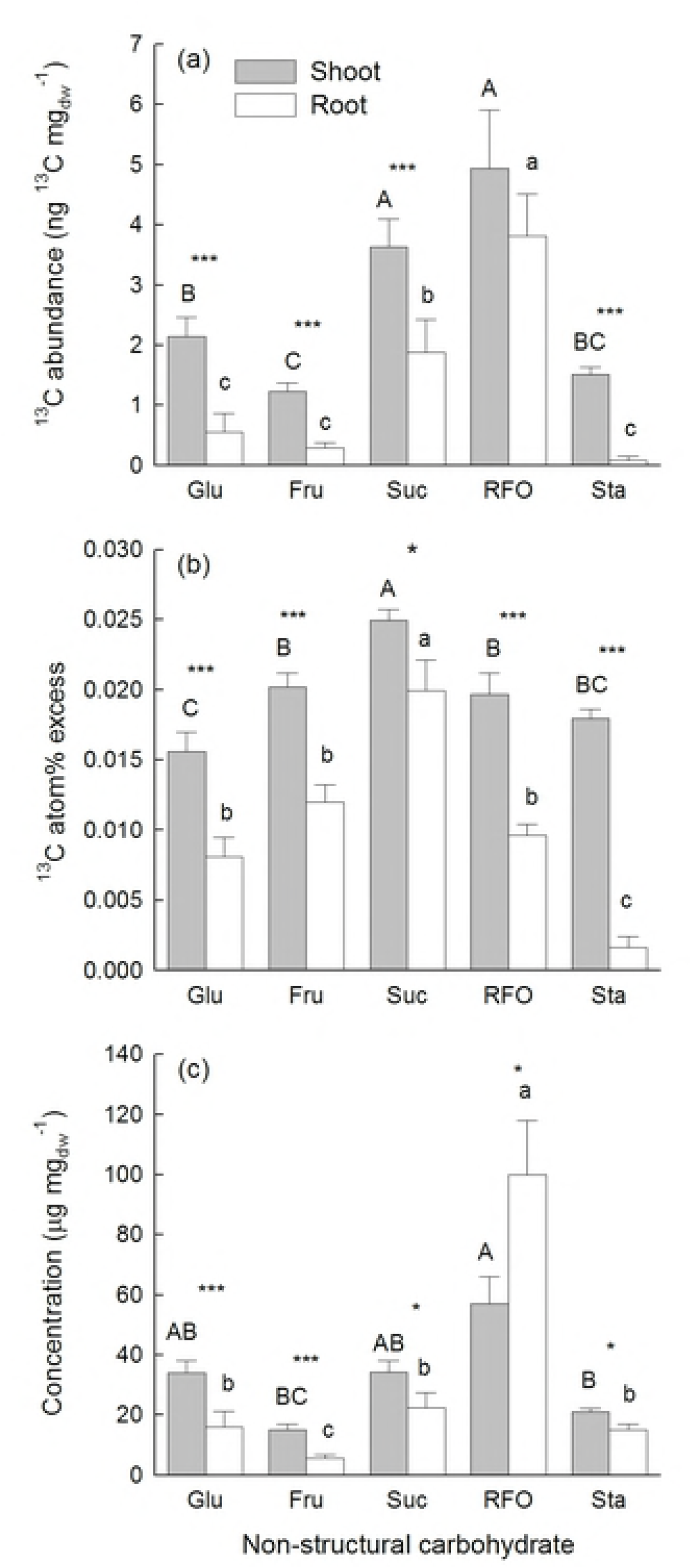
^13^C abundance (a), ^13^C atom% excess (b), and concentration (c) of non-structural carbohydrates (NSC) in shoot and root material. Shown are means (+1 SE) for each studied carbohydrate across all mixtures. Significant differences between roots and shoots are indicated with * P < 0.05, ** P < 0.01, and *** P < 0.001. Results of Tukey’s test applied to test for significant differences among different NSC are marked with upper-case letters for shoots and lower-case letters for roots.

The ^13^C abundance in leaf NSC per unit leaf dry mass was also not different between the 4-species and the 16-species mixtures (S3 Table). The ^13^C abundance was greatest in sucrose (6.97 ± 4.74 ng ^13^C mgdw^-1^) and RFOs (8.45 ± 8.44 ng ^13^C mgdw^-1^) and smaller in starch (2.23 ± 0.70 ng ^13^C mgdw^-1^), glucose (2.66 ± 1.59 ng ^13^C mgdw^-1^) and fructose (1.60 ± 1.11 ng ^13^C mgdw^-1^). The ^13^C abundances in the studied NSC varied to some degree among functional groups (significant interaction NSC-ID × FG-ID; S3 Table, S2 Fig). Especially in grasses, the ^13^C abundance in sucrose and RFOs was considerably higher than in other NSC (Fig 5a). On average, ^13^C atom% excess was lowest in starch and glucose, slightly higher in fructose and RFOs and highest in sucrose (Figs 5e-h). The varying ^13^C abundances in leaf sugars and starch were mainly due to different concentrations of these NSC in leaves of species assigned to different functional groups (Figs 5i-m, S2 Fig).

**Fig 5.**
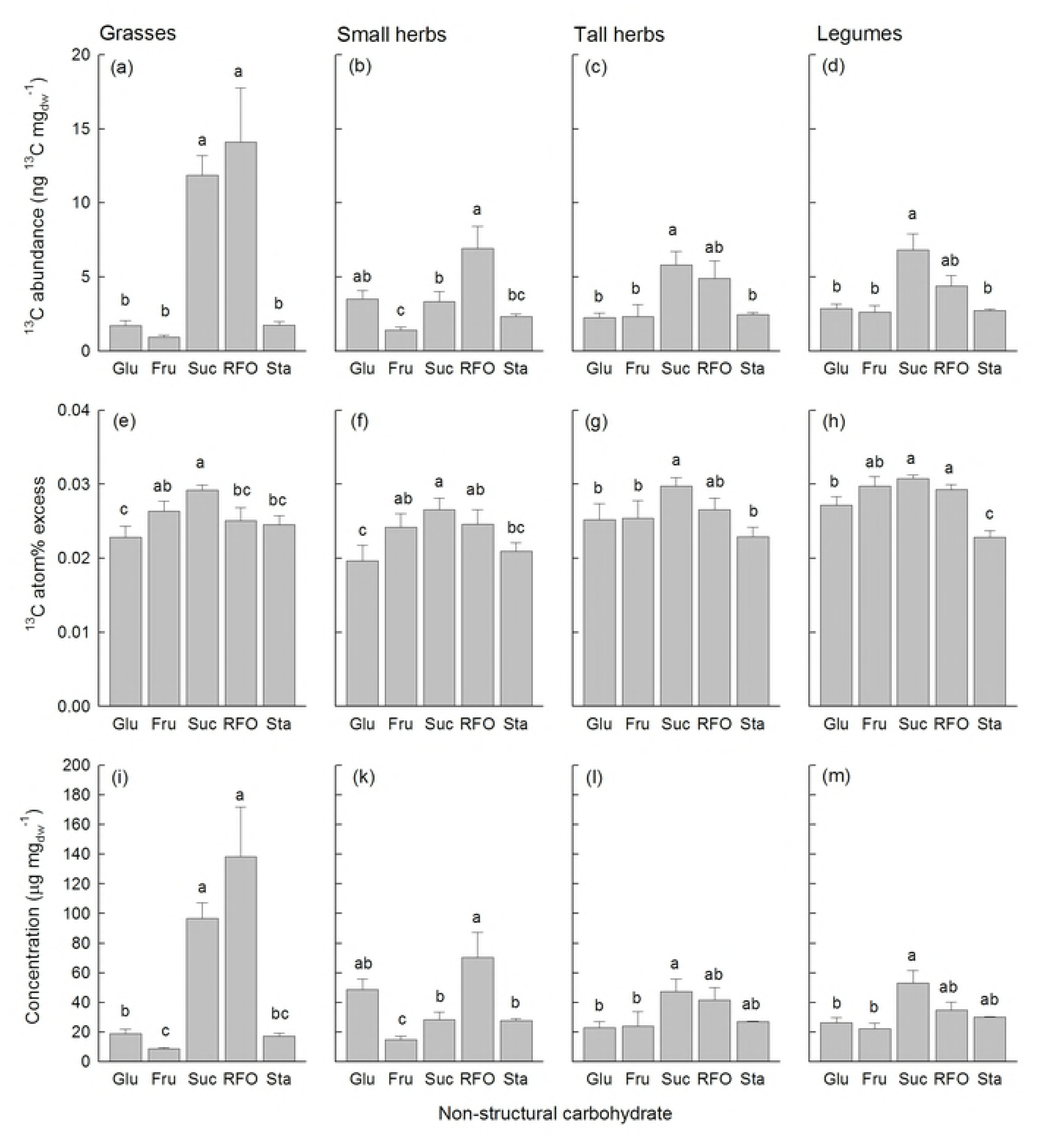
Concentrations and δ^13^C of different non-structural carbohydrates. ^13^C abundances (a-d), ^13^C atom% excess (e-h), and concentrations (i-m) for non-structural carbohydrates in leaves of grasses (a, e, i), small herbs (b, e, k), tall herbs (c, f, l), and legumes (d, h, m). Shown are means (+1 SE) across all mixtures. Results of Tukey’s test applied (P < 0.05) to test for significant differences among non-structural carbohydrates are indicated with letters. Abbreviations are: Glu = glucose, Fru = fructose, Suc = sucrose, RFO = raffinose-family oligosaccharides, Sta = starch.

The atom% excess ^13^C in leaf sugars (glucose, fructose, sucrose, RFO) showed positive relationships with N_Leaf_ and negative relationships with SLA, while the model best explaining variation in ^13^C atom% excess in starch only indicated positive effects of N_Leaf_. The ^13^C atom% excess in RFO was additionally negatively related to LDMC (S4 Table). LDMC was also an important predictor explaining NSC concentrations with negative relationships in glucose, fructose and starch and positive relationships in sucrose and RFO. Consequently, ^13^C abundances in glucose, fructose and starch were negatively related to LDMC, while ^13^C abundances in sucrose showed positive relationships with LDMC. The ^13^C abundance in RFO, however, was best explained by SLA (negative relationships). In addition, N_Leaf_ had positive effects on fructose and negative effects on RFO concentrations and ^13^C abundances (S4 Table).

## Discussion

In our study, we combined species- and community-level measurements of ^13^C abundances in shoot and root biomass as well as in NSC after a three-week lasting ^13^CO_2_ labelling to get insights into the role of plant diversity for carbon gain and allocation.

According to our hypothesis 1, we expected that the amount of assimilated ^13^C (i.e. community-level ^13^C excess) increases with species richness. This hypothesis was only weakly supported by our results because community-level ^13^C excess in shoot and root biomass only tended to be higher in the 16-species than in the 4-species mixtures (Figs 1d and e). Our hypothesis was based on results of many biodiversity experiments including the Jena Experiment, which have shown positive relationships between sown species richness and shoot and root biomass [3]. Our Ecotron experiment was restricted to a subset of the experimental communities of the Jena Experiment, and these communities only displayed a tendency for increased shoot- and root biomass in the studied 16-species compared to the 4-species mixtures (Figs 1g and h). Therefore, we cannot exclude that the chance to observe positive effects of species richness on community-level ^13^C excess was limited by the number of replicates, which we could study in our Ecotron experiment. Nevertheless, Milcu et al. [27] showed in the same Ecotron experiment that net ecosystem C fluxes (i.e. net ecosystem CO_2_ exchange (NEE)) were significantly larger in the 16-species than in the 4-species mixtures. These previous results [27] were based on four observation dates distributed over the regrowth periods after first mowing. However, the diversity effects were strongest in the later phase of the regrowth period when the canopy was well established, which also coincides in time with the ^13^CO_2_ labelling experiment. One possible reason for the stronger diversity effects observed in the previous study [27] is that their measurements referred to the whole area of a monolith, while aboveground biomass for our measurements was harvested on a subplot of 0.495 m^2^ and belowground biomass was determined from soil scores which only represented a small proportion of the monolith. The precision of biomass measurements has a high impact on the determination of ^13^C excess and might therefore have affected our results. One further caveat when comparing the results obtained by Milcu et al. [27] and the present study is three-week duration of the CO_2_ labelling experiment suggesting that the observed patterns did not only represent gross uptake, but most likely some C recycling occurred during this period. We also cannot exclude that a greater loss of ^13^C from plants via higher root exudation rates [45], higher respiration of non- or low-labelled carbon pools [27] or a higher uptake of non-labelled CO_2_ from heterotrophic (and thus non-labelled) soil respiration in the 16-species than in the 4-species mixtures mitigated differences in the ^13^C excess between communities of different diversity.

Community-level shoot ^13^C excess, however, was positively related to canopy leaf N (Fig 2b). Canopy photosynthesis does not only depend on the distribution of N in the canopy, but also on the total amount of leaf area (= leaf area index) of the stand [7]. Increasing leaf area is positively related to the amount of captured light and thus photosynthesis. Analyses of the relative importance of the three variables composing canopy leaf N (g N m^-2^) clearly showed that total leaf area was of prominent importance compared to N_Leaf_ and SLA for community-level shoot ^13^C excess. This result is in line with our hypothesis 1 and previous results [27] showing that higher leaf area index (LAI) and diversity in leaf nitrogen concentrations (FD-N_Leaf_) were important predictors for the positive diversity effects on net carbon fluxes (i.e. NEE, gross ecosystem productivity (GPP)) and relevant variables of carbon uptake efficiency (water use efficiency (WUE), nitrogen use efficiency (NUE)).

As expected (hypothesis 2) shoot ^13^C abundances did not vary between the 4-species and the 16-species mixtures, but N_Leaf_ was also involved in explaining the variation in ^13^C abundances in shoot biomass (Fig 2a). CWM of N_Leaf_ was the most important predictor variable for shoot ^13^C abundances (S2 Table). These results indicated that higher total amounts of N_Leaf_ present in the plant canopy and not many different species-level values of N_Leaf_ (i.e. high FD-N_Leaf_) determine carbon gain integrated over longer time periods. Light absorption per unit N do not differ between dominant tall species and subordinate small species in herbaceous vegetation [46]. The efficiency in acquiring light is a trade-off between tall growth and placing leaves in the uppermost layers to get a high amount of incoming radiation, thereby investing a larger proportion of biomass into supporting tissue consequently reducing efficiency. On the contrary, small-statured species remain in the shade, but achieve a higher efficiency of biomass use to capture photons. Thus, structural traits of plant species play a major role in patterns of light distribution. Other studies suggested that leaf physiological traits rather than plant size-related structural differences explain interspecific differences in carbon gain per unit biomass [22]. In accordance with our hypothesis 2, species-level ^13^C abundances also did not differ between the 4-species and the 16-species mixtures. Lattanzi et al. [47] applied a ^13^C steady-state labelling in grasslands and showed that carbon gain per unit shoot mass increased with increasing plant height in small individuals (species), but it became more independent of plant size among large individuals. In our study, we tested shoot height as well as further leaf traits supposed to be related to leaf gas exchange as predictors for species-level shoot ^13^C abundances. The model best explaining variation in species-level ^13^C abundances combined positive effects of N_Leaf_, g_s_ and H_Shoot_. These traits were also as separate predictors significantly related to species-level ^13^C abundances, but a large scatter in these relationships suggested that their contribution varied among species (S1 Fig). For example, legumes had the largest N_Leaf_ (S5 Table), which could explain their high values of species-level ^13^C abundances (Fig 3). On the contrary, grasses showed the smallest values of g_s_ and grew tallest, while the opposite was the case in small herbs (S5 Table), and species of both functional groups had lower ^13^C abundances in shoot biomass than legumes or tall herbs (Fig 3). Thus, photosynthetic assimilation of both functional groups was likely determined by their position in the plant canopy altering light intensity, air humidity and CO_2_ concentrations.

In general, ^13^C abundance in roots after three weeks of continuous ^13^CO_2_ labelling was small. Our grasslands had a large standing root biomass compared to shoot biomass. Obviously, the recently fixed carbon was mainly used for shoot growth. Concerning the incorporation in specific NSC, the ^13^C atom% excess was as expected (hypothesis 3) greatest in sucrose confirming its main origin from new assimilates (Figs 4b, 5e-h). However, the ^13^C atom% excess in other NSC in the shoots as well as in separately analyzed leaves including starch as storage product was only moderately lower than in sucrose suggesting that recently fixed carbon was partitioned between instant demands and short-term storage. Although the ^13^C atom% excess of sucrose in the roots was only slightly lower than in the shoots, especially root starch showed a very low ^13^C atom% excess indicating that storage in the roots was based on “older” carbon. RFOs, however, do not only serve for transport, but may also function as short-term storage compound [37]. RFOs showed particularly high concentrations in the roots (Fig 4c), and its ^13^C atom% excess in the roots was intermediate compared to those of sucrose and starch suggesting that “fast” and “slow” cycling NSC, i.e. two different pools [48], likely contributed to storage in the roots (Fig 4b). Analyses of NSC in leaves did not show differences in ^13^C atom% excess among functional groups (except for ^13^C atom% excess in glucose). Positive relationships between ^13^C atom% excess in NSC and N_Leaf_ and negative relationships with SLA suggested that species with low-irradiance leaves deeper in the canopy with higher SLA and lower N_Leaf_ [13] contained less recently fixed assimilates. The concentrations of NSC and ^13^C abundances showed large differences among functional groups. Most strikingly, the concentrations of transport sugars (sucrose and RFOs, whereby the latter can also serve as short-term storage) were highest in leaves of grasses. The high concentrations of RFOs in the leaves of grasses suggest that possibly also fructans, which are typically present in grasses [37], contributed to this compound group. RFOs and fructans can have partly overlapping peaks in HPLC-IRMS chromatograms (pers. comm. S. Karlowsky), but since the functions of these carbohydrates are similar this uncertainty would not affect our interpretation. One possible explanation for the higher RFO concentrations is that grasses during the labelling experiments did not have flowers and likely invested in vegetative growth and storage, while legumes and forbs were flowering and allocated more carbon to these “sinks”.

## Conclusions

In summary, our study provided evidence that a larger total leaf area and the functional composition of plant communities play a prominent role for increased carbon gain of plant communities. The importance of leaf N in explaining species- and community-level C uptake underscores the close relationships between C and N metabolism. Although the effect of increased species richness on the amount of assimilated carbon was weak in our Ecotron experiment, the greater variety of traits related to photosynthesis and plant positioning in the canopy involved in explaining species-level C uptake highlights that a greater diversity in the expression of plant traits, which is often positively related to species richness, is important for high community-level carbon gain.

## Acknowledgments

We thank W.W. Weisser and A. Ebeling for coordination of the Jena Experiment, the Ecotron team for running the experiment in Montpellier, U. Gerighausen for sample preparation, H. Geilmann and W.A. Brand (Jena) for stable isotope analyses and A. Ackermann (Zurich) for elemental analyses.

## Supporting information

**S1 Fig.** Scatterplots between species-level ^13^C abundance in shoots and functional traits.

**S2 Fig**. Non-structural carbohydrate concentrations and δ^13^C in relation to functional groups.

**S1 Table.** Species composition of studied mixtures.

**S2 Table.** Analysis of community-level shoot ^13^C abundance and ^13^C excess in relation to functional composition.

**S3 Table.** Analysis of leaf non-structural carbohydrate concentrations and δ^13^C in relation to species richness and functional groups.

**S4 Table.** Analysis of leaf non-structural carbohydrate concentrations and δ^13^C in relation to plant traits.

**S5 Table.** Analysis of plant traits in relation to species richness and functional group identity.

**S1 Method.** Extraction and measurement of non-structural carbohydrate concentrations and δ^13^C.

**S2 Method.** Measurement of aboveground plant traits.

